# Polymorphic regions in BA.2.12.1, BA.4 and BA.5 likely implicated in immunological evasion of Omicron subvariant BQ.1.1

**DOI:** 10.1101/2022.11.20.517236

**Authors:** Pierre Teodosio Felix

## Abstract

In this work, 45 Spike glycoprotein Chain B polypeptides were used in the subvariants BA.2.12.1, BA.4 and BA.5 were recovered from GENBANK. All sequences were publicly available on the National Biotechnology Information Center (NCBI) platform. The results indicate the existence of informative polymorphic and parsimony sites that may be implicated in the level of diversity of the studied strains, as well as reflect the immunological evasion potential of the subvariant BQ1.1. of the variant Ômicron d and SARS-CoV-2. The results also suggest the formation of ancestral polymorphism with slight retention, and the probable is responsible the diversity of the whole studied set.

## 1. Introduction

In less than a year, many mutations in the have given rise to new subvariants that compromise vaccine-induced immunity (Panke *et al,* 2022). The initial subvariant of Ômicron has already shown, from the beginning, a strong immunological escape of immunity induced by two doses of the mRNA vaccine, which was only effective with the booster dose (Evans *et al.*, 2022; Gruell *et al.*, 2022). Among the subvariants that emerged, the subvariant BA.2 showed a greater reinfection capacity than BA.1, in addition to a slight increase in immunological evasion (Stegger *et al.*, 2022), succeeded by the subvariants BA.4 and BA.5 with identical S proteins, exhibiting greater immunological escape (Kimura *et al.*, 2022; Tuekprakhon *et al.*, 2022). Although BA.4 and BA.5 are identical in terms of Spike protein mutations, they share and differ in mutations that are outside the Spike protein. These mutations affect viral replication, infection rate, and treatment resistance. Both BA.4 and BA.5 reverse two mutations back to the original virus, Orf6 D61 and NSP4 L438, which replicate the virus by negative regulation of proteins, enzymes and various other signals. The protein residing in NSP4, L438 NSP4, is involved in the formation of a double membrane vesicle, which also potentiates viral replication (CAS, 2022). The subvariant BA.4/5, in turn, generated greater variation in SARS-CoV-2, causing strains such as BQ.1.1, currently responsible for the largest contamination frequency according to centers for disease control and prevention, 2022.

Trying to understand the diversity of the immunological evasion potential present in the BQ.1.1 strain, we of the laboratory team of the Laboratory of Population Genetics and Computational Evolutionary Biology (LaBECom-UNIVISA), performed a phylogenetic work with 45 polypeptides of chain B of glycoprotein Spike in the subvariants BA.2.12.1, BA.4 and BA.5, available in the database of the National Biotechnology Information Center (NCBI).

## 2. Methodology

### 2.1. Data Bank

45 Spike Glycoprotein Chain B polypeptides in the BA.2.12.1, BA.4 and BA.5 subvariants *were* retrieved from GENBANK (molecule 7XNS, chain B, release Aug 31, 2022; deposition: Apr 29, 2022; class: VIRAL PROTEIN; source: Mmdb_id: 217871, Pdb_id 1: 7XNS; Exp. method: Electron Microscopy) and that were used in the work of Cao *et al,* 2022. Theseviral proteins contained 1273 aa that were aligned using the MEGA X program (Kumar *et al.*, 2018) and ambiguous data, lost data and gaps were excluded, generating a content of 987 analyzable sites.

### 2.2. Inferred Ancestral Sequencies

Analyses: For the measurements of the ancestral sequences, the Maximum Likelihood method with Bootstrap of 100 pseudo-replications was used and with the Jones-Taylor-Thornton Aminoacid Substitution (JTT) model. A gamma distribution with invariant sites (G+I) was used. All gaps were extracted and tree was automatically generated by the Standard heuristic method - NJ/BioNJ.

### 2.3. For viewing variable sites

The graphical representation of the sites was made using the WEBLOGO v3 software, described by CROOKS *et al.*, 2004.

## 4. Results

### 4.1. General properties of the 45 segments of Chain B of spike glycoprotein

The 45 viral proteins were recovered from GENBANK on November 17, 2022. These haplotypes contained 1273minoacids that, after being aligned using the MEGA X program (Kumar *et al.*, 2018), generated a segment with 987 conserved sites and 286 polymorphic sites. In total, all 244 sites were informative parsimonium sparpmium, which, once treated for the exclusion of lost sites, ambiguous sites and gaps, generated a segment of 73 analyzeable sites. The graphic representation of these sites could be seen in a logo built with the PROGRAM WEBLOGO 3. (CROOKS *et al.*, 2004), where the size of each nucleotide. proportional to its frequency for certain sites (Figure 1 and Figure 2).

**Figure 1:**
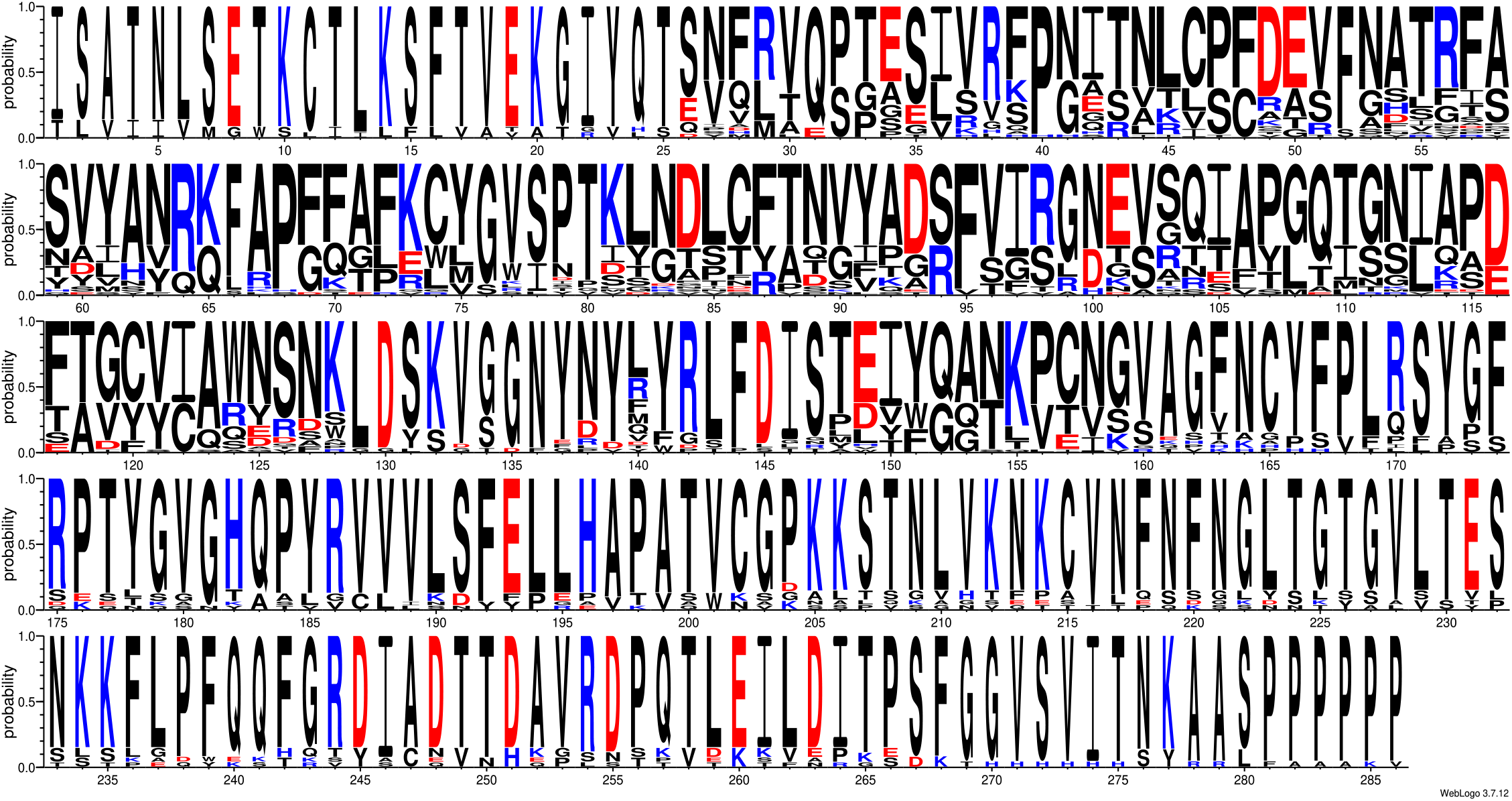
Graphic representation of the 286 polymorphic sites of the 45 sequences of Chain B of Spike glycoprotein in subvariants BA.2.12.1, BA.4 and BA.5, retrieved from GENBANK.

**Figure 2:**
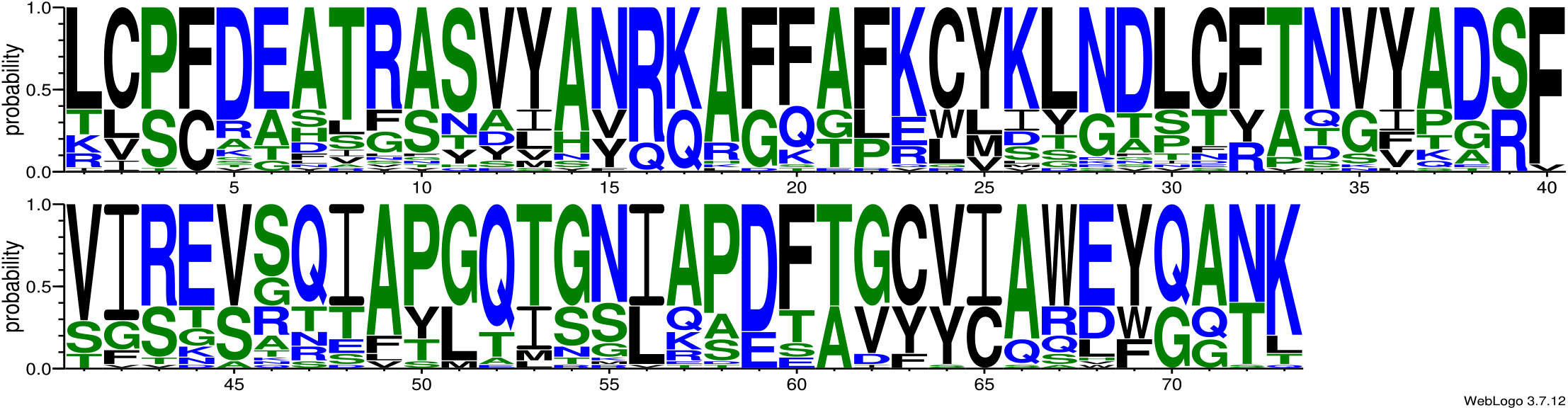
Graphic representation of 73 parsimonium-informative sites of the 45 sequences of Spike glycoprotein Chain B in the subvariants BA.2.12.1, BA.4 and BA.5, retrieved from GENBANK.

Using the UPGMA method for the 73 parsimony-informative sites, it was possible to understand that the 45 haplotypes comprised five distinct groups, with reasonable haplotypic sharing. The maximum likelihood maps found the presence of phylogenetic signal for all data sets, presenting more than 70% of the quartets resolved (data not shown). The evolution model GTR+I+G was the one that best represented the differences between groups, with a *significant bootstrap value* (65%) supporting clade separation (Figure 3).

**Figure 3:**
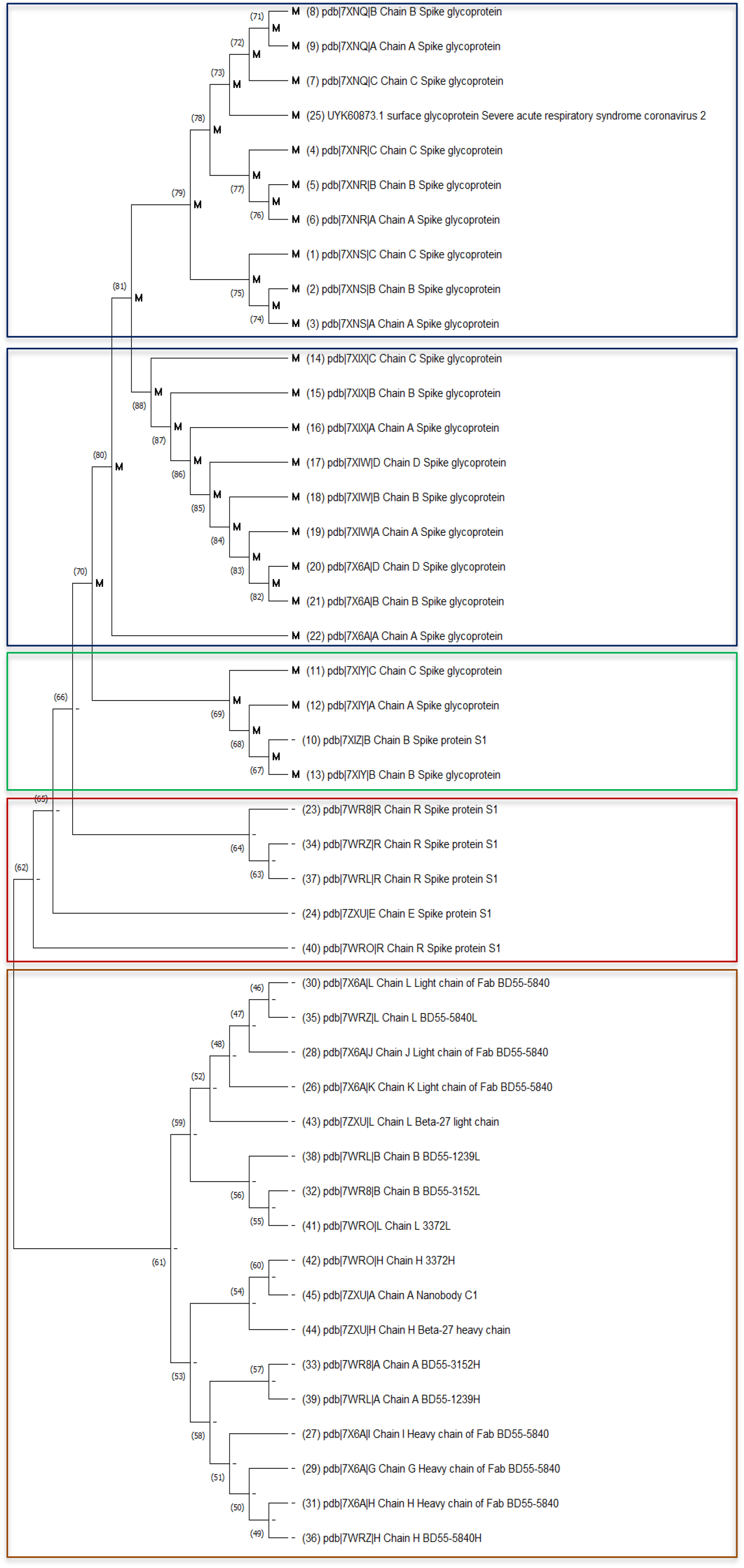
Inferred Ancestral Sequences. Ancestral states were inferred using the Maximum Likelihood method [1] and JTT matrix-based model [2]. The tree shows a set of possible amino acids (states) at each ancestor node based on their inferred likelihood at site 1. For each node only the most probable state is shown. The initial tree was inferred using the method. The rates among sites were treated as a Gamma distribution with invariant sites using 5 Gamma Categories (Gamma with invariant sites option). This analysis involved 45 amino acid sequences. There was a total of 73 positions in the final dataset. Evolutionary analyses were conducted in MEGA X [3]

## 5. Discussion

The polypeptides studied did not present significant level of structuring perhaps because they did not have considerable inter-haplotipica variation, these being the main components in moderate structuring. These data suggest that the absence of a degree of structuring is related to a probable loss of intermediate haplotypes in the course of mutations.

The levels of structuring presented here were supported by simple methodologies of phylogenetic pairing such as UPGMA, revealing a continuous pattern of genetic divergence (a fact that does not support the occurrence of geographical isolationstemming from past events), was observed a reasonable number of branches with many clear mutational steps, which is fixed accompanying the dispersal behavior and/or loss of intermediate haplotypes.

The entire studied group showed a significant polymorphic pattern with subtle differences in its internal haplotypic sharing (parsimonious-informative sites). The search for inter-haplotypic variations were attempted to hierarchize the covariance components by their intra- and inter-individual differences,generating a pattern that supports the idea that the significant differences found for this group were shared more in their number than in their form, once again. that the result of the estimates of the mean evolutionary divergence found within and between them were not significant. Analyzing all the polymorphisms found, it can be suggested that the variation in protein products for the Spike protein should be considered, since these evidence rapid and non-silent mutations that significantly increase the genetic variability sars-cov-2 virus in a very short time, and may be one of the factors that elect the subvariants BA.2.12.1, BA.4 and BA.5 of Ômicron, as well as the recent BQ1.1., as highly specialized in the current immunological evasion observed.

## References

Cao, Y., Yisimayi, A., Jian, F. et al. BA.2.12.1, BA.4 and BA.5 escape antibodies caused by Omicron infection. Nature 608, 593–602 (2022). https://doi.org/10.1038/s41586-022-04980-y.

Centers for Disease Control and Prevention (2022). COVID Data Tracker. Atlanta, GA: US Department of Health and Human Services, CDC; 2022.

Chemical American Society (2022). Why the BA.5 variant of Ômicron avoids vaccines. Available in: https://www.cas.org/pt-br/resources/blog/covid-omicron-ba5-variant. Accessed 11/20/2022.

Crooks GE, Hon G, Chandonia JM, Brenner SE WebLogo: A sequence logo generator, Genome Research, 14:1188–1190, (2004) [Full Text].

Evans, J.P., Zeng, C., Qu, P., Faraone, J., Zheng, Y.M., Carlin, C., Bednash, J.S., Zhou, T., Lozanski, G., Mallampalli, R., et al. (2022). Neutralization of SARS-CoV-2 Ômicron sub-lineages BA.1, BA.1.1, and BA.2. Cell Host Microbe 30, 1093–1102.e1093.

Gruell, H., Vanshylla, K., Tober-Lau, P., Hillus, D., Schommers, P., Lehmann, C., Kurth, F., Sander, L.E., and Klein, F. (2022). mRNA booster immunization elicits potent neutralizing serum activity against the SARS-CoV-2 Omicron variant. Nat Med 28, 477–480.

Kimura, I., Yamasoba, D., Tamura, T., Nao, N., Suzuki, T., Oda, Y., Mitoma, S., Ito, J., Nasser, H., Zahradnik, J., et al. (2022). Virological characteristics of the SARS-CoV-2 Omicron BA.2 subvariants, including BA.4 and BA.5. Cell 185, 3992–4007.e3916.

Kumar, S., Stecher, G., Li, M., Knyaz, C., & Tamura, K. (2018). MEGA X: Molecular evolutionary genetics analysis across computing platforms. Molecular Biology and Evolution, 35(6), 1547–1549. https://doi.org/10.1093/molbev/msy096.

Panke Qu, John P. Evans, Julia Faraone, Yi-Min Zheng, Claire Carlin, Mirela Anghelina, Patrick Stevens, Soledad Fernandez, Daniel Jones, Gerard Lozanski, Ashish Panchal, Linda J. Saif, Eugene M. Oltz, Kai Xu, Richard J. Gumina, Shan-Lu Liu (2022). Distinct Neutralizing Antibody Escape of SARS-CoV-2 Omicron Subvariants BQ.1, BQ.1.1, BA. 4.6, BF.7 and BA.2.75.2. bioRxiv 2022.10.19.512891; doi: https://doi.org/10.1101/2022.10.19.512891.

Stegger, M., Edslev, S.M., Sieber, R.N., Ingham, A.C., Ng, K.L., Tang, M.-H.E., Alexandersen, S., Fonager, J., Legarth, R., Utko, M., et al. (2022). Occurrence and significance of Omicron BA. 1 infection followed by BA. 2 reinfection. medRxiv.

Tuekprakhon, A., Nutalai, R., Dijokaite-Guraliuc, A., Zhou, D., Ginn, H.M., Selvaraj, M., Liu, C., Mentzer, A.J., Supasa, P., Duyvesteyn, H.M.E., et al. (2022). Antibody escape of SARS-CoV-2 Omicron BA.4 and BA.5 from vaccine and BA.1 serum. Cell 185, 2422–2433.e2413.

